# Repeated drought amplifies DNA methylation at shared transposable element targets in *Quercus lobata*

**DOI:** 10.64898/2026.07.14.738593

**Authors:** L. D. Peck, V. L. Sork

## Abstract

Long-lived trees experience repeated environmental stress over their lifetimes, and droughts are becoming more frequent with climate change. Environmentally responsive DNA methylation is widespread in plants, but how it develops over repeated stress, and where in the genome it occurs, is poorly understood in trees. We generated Nanopore methylomes for valley oak (*Quercus lobata)* seedlings sampled before, during and after repeated drought, alongside well-watered controls. Repeated drought was associated with widespread gains in CHH methylation, with drought-enhanced gains in 14,055 regions, compared with 26 regions showing the reverse. Most of the drought-specific excess accumulated after the first drought. Gains were concentrated in transposable elements, particularly small, high-copy DNA transposons near genes and in chromatin that was accessible before treatment. Drought and well-watered plants gained methylation at the same transposable element (TE) copies more frequently than expected from their genomic characteristics, revealing a partly shared targeting landscape. CHH gains occurred near drought-induced genes involved in abscisic acid signalling and dehydration responses, but were not generally associated with transcriptional induction. We detected no increase in TE insertions under drought. Our findings suggest that environmental stress amplifies CHH methylation within an existing, copy-specific targeting landscape shaped by genomic and chromatin characteristics, providing new insights into how long-lived plants respond epigenetically to recurrent environmental stress.

## Introduction

Trees experience decades to centuries of environmental change over their long lives. Unlike annual plants or short-lived perennials, which can adapt to environmental change over relatively short generation times through natural selection, trees must repeatedly adjust gene expression and physiological responses to changing conditions within a single lifespan (Bräutigam *et al*., 2013; Aitken & Whitlock, 2013). How these repeated molecular responses are coordinated remains poorly understood. Moreover, because climate change is predicted to increase the frequence and severity of droughts (Urban, 2015), it is important to understand the molecular mechanisms that enable trees to respond to repeated drought events.

In plants, physiological responses to environmental stress are coordinated through changes in gene expression (Nicotra *et al*., 2010; Takahashi *et al*., 2020; Zhang *et al*., 2022). These transcriptional responses are determined not only by promoter sequences but also by chromatin state and the broader genomic context surrounding genes (Berger, 2007; Ricci *et al*., 2019; Marand *et al*., 2021; Pozzi *et al*., 2025). Gene-proximal transposable elements (TEs) and DNA methylation are increasingly recognized as important components of this regulatory landscape (Law & Jacobsen, 2010; Niederhuth & Schmitz, 2017; Klein & Anderson, 2022). Although methylation occurs in CG, CHG, and CHH sequence contexts (where H=A, T, or C), CHH methylation is particularly dynamic because it depends upon continual targeting through the RNA-directed methylation pathway (Chan *et al*., 2006; Cokus *et al*., 2008; Secco *et al*., 2015; Dubin *et al*., 2015; Rajkumar *et al*., 2020). Consequently, CHH methylation has emerged as one of the most environmentally responsive components of the plant methylome (Sork *et al*., 2026; Sanchez-Lucas *et al*., 2026) and is enriched at TEs where it may influence nearby transcription while maintaining TE repression (Law & Jacobsen, 2010; Gent *et al*., 2013). Studies in trees have documented drought-associated methylation changes (Xu *et al*., 2018; Wang *et al*., 2023; Zhou *et al*., 2023; Peña-Ponton *et al*., 2024), and other plant studies have implicated transposable elements in shaping stress-responsive transcription (Srikant & Drost, 2021). Recent comparative analyses in two *Quercus* species suggest that CHH methylation is environmentally induced and associated with TE distribution (Sork *et al*., 2026; Sanchez-Lucas *et al*., 2026), as well as associated with gene expression in *Q. lobata* (Sork *et al*., 2026). However, whether repeated drought exposure alters methylation at shared TE targets, and how these changes relate to gene expression, remains poorly understood (Peck & Sork, 2024).

In this study, we investigated how environmental stress affects the regulatory landscape in a tractable study system, valley oak seedlings (*Q. lobata* Née). Using long-read genomes and DNA methylomes collected across repeated drought and well-watered treatments, along with a subset of individuals with transcriptomes and chromatin accessibility, we tested whether (i) repeated drought induces CHH methylation, and if so, at which genome regions; (ii) these methylation patterns are associated with gene expression; and (iii) repeated drought redirects or amplifies methylation targeting. Our results show that repeated drought amplifies CHH methylation at specific, shared genomic TE targets without evidence of increased TE activity. Whether these methylation changes contribute to transcriptional responses remains an important question for future investigation.

## Results

### Repeated drought produces a delayed amplification of CHH methylation

Our observations of *Quercus lobata* seedlings from seven maternal trees (hereafter referred to as families, Fig. S1A) growing under drought and well-watered treatments revealed increasing differences in physiological measures over three time points: pre- drought (T1); after 12 days of drought (T2); and following recovery and a second 12-day drought (T3; Fig. 1A and Fig. S2). We sequenced eight seedlings from two families (24 methylomes in total) and found that repeated drought amplified widespread gains in CHH methylation. Between T1 and T3, 14,055 regions met our stringent criteria for showing greater methylation gains in all four drought-treated plants relative to well-watered controls, compared with only 26 regions showing the reciprocal pattern (Fig. 1B). Almost all CHH changes were gains; CHG regions were fewer (357) and predominantly losses, and few CG regions changed. A plant-blocked DSS pipeline revealed the samepattern (see Supplementary Results).

**Figure 1.**
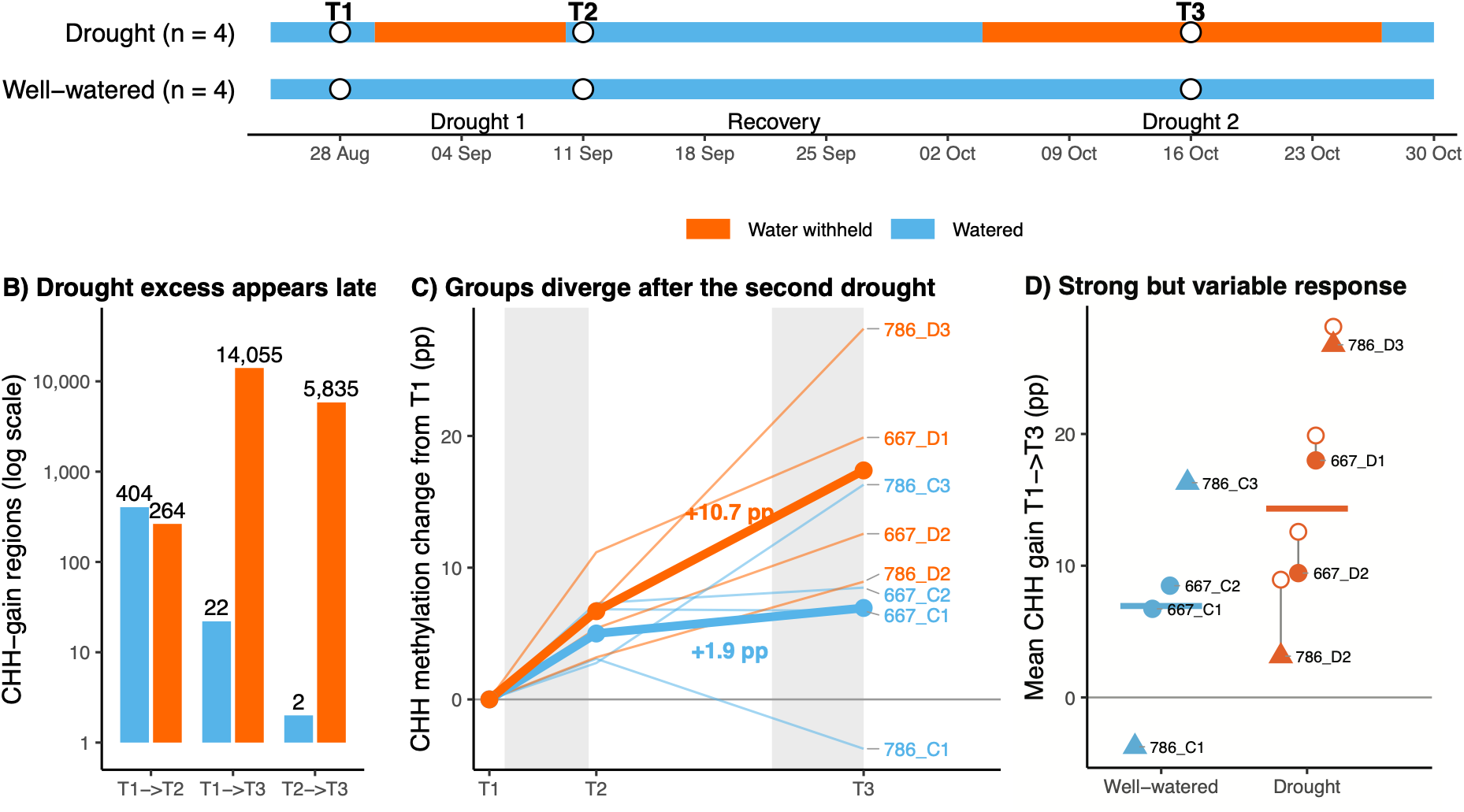
Repeated drought amplifies CHH methylation in the second drought stress. **(A)** Design: valley oak seedlings (two maternal families) were sampled before treatment (T1), after a first 10-day drought (T2, 11 Sep 2023), and after 24 days of recovery, 10 days into a second drought (T3, 16 Oct; water was withheld 4–27 Oct); drought and well-watered, n = 4 plants each. **(B)** Number of CHH-gain regions (log scale) in drought and well-watered plants for each interval (pooled-read Fisher test, all four plants gain ≥ 5 pp, read-level drought × time interaction and plant-mean difference ≥ 5 pp). **(C)** Mean CHH change from T1 at the drought T1-T3 gain regions; thin lines, individual plants; thick lines, group means; grey, drought periods. Regions were selected on T1 and T3 only, so T2 is an independent measurement. **(D)** Mean T1-T3 CHH gain per plant (one point per plant; shape, family). Drought plants (filled) are measured at regions identified from the other three drought plants, so no plant’s own reads were used to select its regions; open circles give the same plant’s gain at the full set of regions in C). Well-watered plants, full set. Bars, group means (Well-watered 6.9 pp; Drought 14.3 pp).

The drought-associated excess emerged primarily after the first drought treatment. Between T1 and T2, more regions were identified in controls than drought-treated plants (404 vs 264). At regions subsequently identified as drought-enhanced between T1 and T3, both groups showed modest early gains (mean ΔCHH = 0.067 and 0.050, respectively), but only drought-treated plants continued to gain substantially between T2 and T3 (+0.107 vs +0.019; Fig. 1C). Approximately 84% of the excess gain therefore accumulated during this later interval. Gains were spatially restricted, with little corresponding change in neighbouring regions within 1 kb. Although the magnitude of CHH methylation gain varied considerably among drought-treated individuals, each showed gains at regions identified without its own sequencing data in a leave-one-out analysis (Fig. 1D).

### CHH gains target specific transposable elements shared between drought and control plants

Drought-enhanced CHH methylation gains preferentially occurred in transposable elements (TEs), particularly small, high-copy DNA transposons near genes and in initially accessible chromatin. After accounting for baseline CHH methylation and sequencing depth, TE-containing regions had 2.5-3-fold higher odds to gain methylation than non-TE regions. Enrichment was strongly dependent on TE class. Relative to detection-matched regions, CHH gain was enriched on PIF-Harbinger (2.5 fold), DNA unclassified (2.4 fold), hAT-Tip100 (2.4), and hAT-Ac (1.9), and depleted on LTR/Copia and LTR/Ty-3 (formerly Gypsy (Wei *et al*., 2022)) elements. Relative to regions matched for initial methylation and sequencing depth, gains were enriched 1.9-fold upstream and 2.0-fold downstream of genes (within 1.5 kb), and 3.0-fold within 500 bp of a T1 ATAC-seq peak, or accessible chromatin region (Figure 2B). These associations persisted after adjustment for TE class, length, copy number, and gene proximity (upstream odds ratio = 1.50; downstream = 1.75; accessible chromatin = 2.01). Gains preferentially affected shorter elements (odds ratio = 0.48 per standard deviation of length), and higher-copy elements (odds ratio = 1.56 per SD).

**Figure 2.**
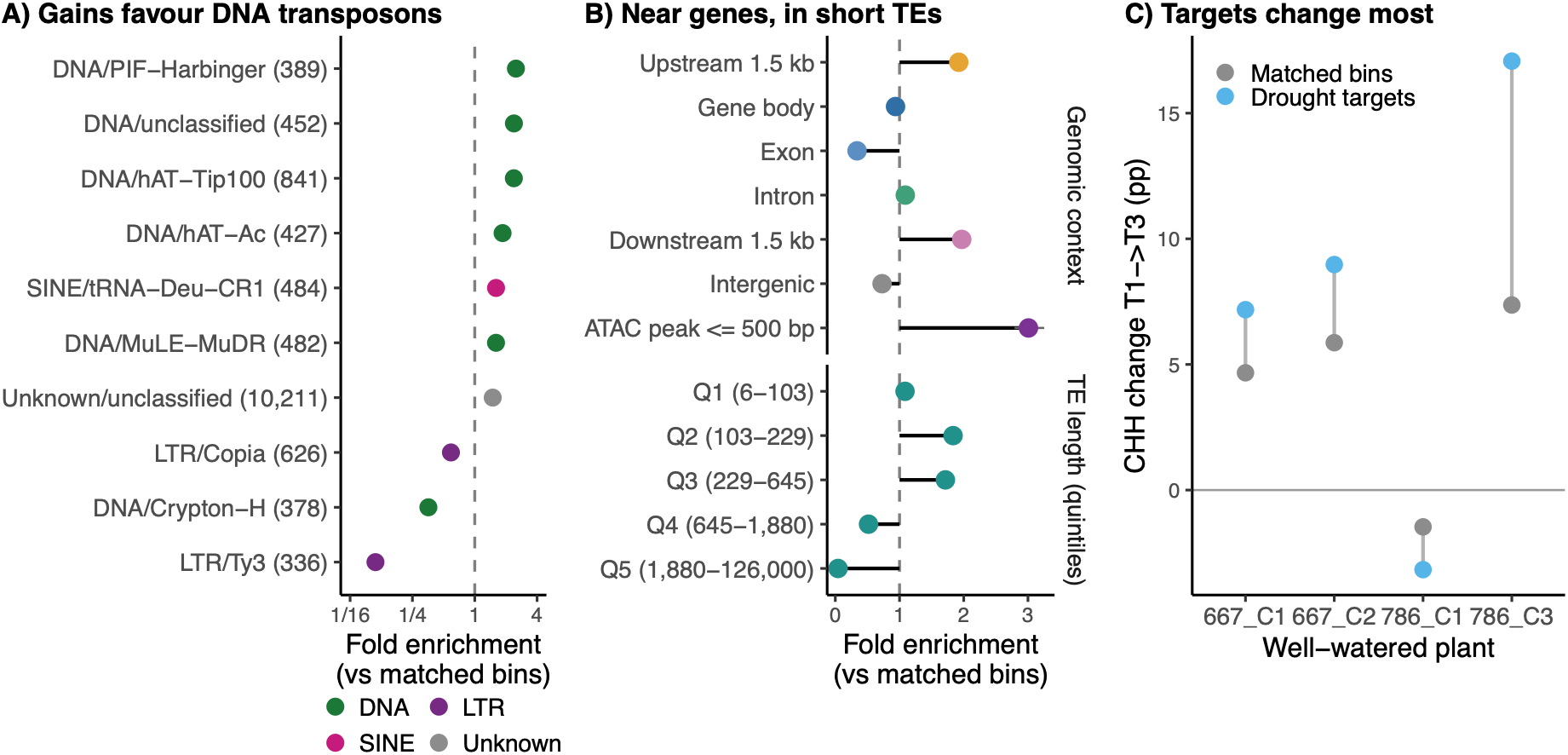
Drought-associated CHH methylation gains preferentially target short, gene-proximal transposable elements (TEs), with partial overlap between treatments. **(A, B)** Fold enrichment of 200-bp bins within drought-enhanced T1–T3 CHH-gain regions relative to tested bins matched on baseline (T1) CHH methylation decile and sequencing-depth quintile. A value of 1 indicates no enrichment relative to matched bins; error bars show 95% confidence intervals from a 1-Mb genomic block bootstrap. **(A)** Enrichment across TE categories with >100 gain bins, coloured by TE order (log2 scale; numbers in parentheses indicate gain-bin counts). **(B)** Enrichment by genomic context, proximity to a pre-drought (T1) ATAC-seq peak (≤500 bp), and TE length quintile (among TE-overlapping bins; linear scale). Exons and introns are subsets of gene bodies. Enrichments in A and B were estimated separately; jointly adjusted effects are shown in Fig. S2e. **(C)** T1–T3 CHH methylation changes in individual well-watered plants at drought-gain TE bins (blue) and matched non-gaining TE bins (grey), matched on baseline methylation, sequencing depth, TE category, genic context and accessibility. Baseline methylation deciles were calculated from the other three well-watered plants, excluding the focal plant, and drought-gain bins were identified without control-plant reads.

Importantly, drought did not establish a distinct methylation target landscape. Regions gaining CHH methylation in well-watered controls showed similar genomic characteristics, including enrichment near genes and within accessible chromatin (Fig. S3). Moreover, drought and control gains occurred at the same TE copies more frequently than expected based on their measured genomic characteristics. Over T1-T3, shared gains showed 2.3 times the expected overlap after adjustment (Table S2). Consistent with this shared targeting, three of four well-watered plants gained more CHH methylation at drought-targeted TE regions than at matched regions, although the fourth showed greater methylation loss at drought targets (Fig. 2C). Thus, repeated drought amplified CHH methylation within a genomic targeting landscape partly shared with well-watered controls, including preferential gains at some of the same TE copies.

### CHH gains occur near drought-responsive genes without increased transposable element mobilisation

CHH gains near genes showed limited associations with drought-responsive transcription. Genes with upstream CHH-gaining TEs were modestly more likely to be induced (odds ratio = 1.22, nominal *P* = 0.018) and were no more likely to be repressed (odds ratio = 0.98, Table S3). Several drought-induced genes involved in abscisic acid (ABA) signalling and dehydration responses, including *COR47, AFP3, ABF2/AREB1, DREB2A,* and *CIPK6*, carried upstream CHH-gaining TEs (Table S4). Among induced genes, ABA-response genes showed a tendency towards greater odds of upstream CHH gain after accounting for differences in detection opportunity, which was consistent across alternative model specifications although not statistically significant (odds ratio = 1.74, *P* = 0.14, Table S3). Thus, we observed a weak association between upstream CHH gain and transcriptional induction, but no compelling evidence of a specific regulatory relationship.

Despite extensive methylation gains at TEs, we found no clear evidence of increased TE mobilisation under drought. Per-leaf rates of candidate new TE insertions closely tracked those of non-TE insertions, which provided an estimate of library-specific artefact background (single read calls, r = 0.98 across 24 leaves; Fig. S5). Leaves formed during the experiment showed no drought-specific increase relative to this background (rate ratio = 1.01; Fig. S5B, Table S5). Multi-read insertions retained after cross-leaf re-genotyping were rare (231 across 24 leaves) and occurred at low within-leaf allele fractions (median = 0.05 in both treatments). The estimated drought effect on multi-read insertion rates was highly uncertain (rate ratio = 2.3; Table S6). Therefore, extensive CHH gains at resident TEs were not accompanied by a detectable increase in new insertions, although our ability to resolve changes in rare insertion events was limited.

## Discussion

Our study reveals that repeated drought amplifies CHH methylation at a shared set of transposable elements (TEs) in *Quercus lobata*. The largest methylation gains occurred following the second drought, but these changes were not restricted to drought-treated plants: well-watered controls also gained methylation at the same TE copies more frequently than expected based on their genomic features. This finding suggests that drought enhances methylation at an existing set of genomic targets rather than establishing an entirely distinct targeting landscape. The predominance of CHH hypermethylation is consistent with environmentally responsive methylation pathways reported in other plants (Secco *et al*., 2015; Rajkumar *et al*., 2020; Zhou *et al*., 2023; Peña-Ponton *et al*., 2024; Sork *et al*., 2026; Zhang *et al*., 2026). However, our temporal sampling cannot distinguish between methylation maintained during recovery and renewed or cumulative methylation gains following the second drought. Thus, although repeated drought was associated with a larger epigenomic response, we cannot establish whether this trend represents priming or epigenetic stress memory.

The genomic distribution of CHH gains provides insight into how these targets may be selected. Gains preferentially affected smaller, high-copy DNA transposons, particularly those located near genes and in regions of chromatin accessible before drought. The association with accessibility persisted after accounting for TE identity, length, copy number, and genomic position, suggesting that pre-existing chromatin context contributes to the fact that particular TEs gain methylation. Indeed, this interpretation is consistent with evidence that TE identity, age, and genomic position influence methylation dynamics and regulatory potential (Hollister & Gaut, 2009; Hollister *et al*., 2011; vonHoldt *et al*., 2012; Choi & Purugganan, 2018; Wang & Baulcombe, 2020). Together with the enrichment of shared gains at individual TE copies, these findings are consistent with the preferential targeting of a subset of gene-proximal elements, potentially through RNA-directed DNA methylation (RdDM) (Chan *et al*., 2006; Cokus *et al*., 2008). Such elements occupy an important interface between TE silencing and gene regulation (Hollister & Gaut, 2009; Gent *et al*., 2013; Klein & Anderson, 2022). Nevertheless, neither RdDM activity nor drought-induced changes in chromatin accessibility were measured directly, and additional copy-specific features or technical effects may contribute to the observed targeting patterns.

Despite the proximity of CHH gains to genes, we found no statistically significant evidence that they were associated specifically with transcriptional induction. Several drought-responsive genes involved in abscisic acid signalling and dehydration responses carried upstream CHH-gaining TEs, but this association was not significant after accounting for gene induction. Furthermore, we detected no increase in candidate TE insertions under drought, suggesting that the methylation response primarily affected resident elements rather than accompanying detectable TE mobilisation. This study identifies a structured epigenomic response to repeated environmental stress, providing a foundation for future investigations of its underlying mechanisms and potential regulatory consequences.

## Materials and Methods

### Sample design and drought assays

In October 2022, acorns were sampled from seven *Q. lobata* trees growing in oak savannah habitat growing in Sedgwick Reserve, located in the Santa Ynez Valley, Santa Barbara County, CA, (Figure S1) and administered by the University of California Santa Barbara, as part of the University of California’s natural reserve system.

To assess the physiological responses to the soil drying and well-watered treatments, we monitored the progress of all seedlings from each maternal tree (Tree IDs: 530, 667, 27, 165, 786, 186, 896). Acorns were surface-sterilized and planted in a randomized design in the greenhouse at UCLA. In July 2023, after approximately seven months since germination, seedlings from each maternal family (i.e., half-siblings from seven maternal trees) were randomly assigned to drought and well-watered treatment groups. Twice weekly, seedling transpiration was measured using a Licor Li-600 porometer and soil moisture measured using a Hobo MX2307 probe to monitor seedling response to drought. Three readings were taken per seedling from the newest fully expanded leaves. Well-watered seedlings were watered every few days, while drought seedlings underwent a first drought period for twelve days, after which they were watered until their stomatal conductance rate returned to the same level as the well-watered seedlings. A second and longer drought period for 23 days was commenced. We followed methods developed in previous drought studies completed (Gugger *et al*., 2017; Mead *et al*., 2019; Steele *et al*., 2025).

For the genome-wide methylation profiling, we selected eight half-sib seedlings from two maternal families (667 and 786), comprising two seedlings per family in each treatment (drought and well-watered). Leaf samples were collected before treatment application (time point 1, T1); at the end of the first twelve-day treatment (time point 2, T2); and twelve days into the second drought treatment following rewatering (time point 3, T3). Each seedling was sampled across all three time points, with drought responses evaluated relative to two complementary controls: (i) contemporaneous well-watered seedlings from the same maternal families; and (ii) pre-treatment (T1) measurements from the same individual seedlings. This repeated-measures design allowed drought-associated changes to be assessed relative to baseline variation and contemporaneous controls.

### ATAC-seq, peak calling and characterisation of accessible chromatin regions

Chromatin accessibility was profiled using ATAC-seq in T1 leaves of two drought-assigned seedlings (786_D1 and 786_D3), sampled before treatment. Nuclei were isolated, purified, and subjected to Tn5 tags, and ATAC-seq libraries were prepared following the protocol described by Minow *et al*. (2025). Libraries were sequenced to generate paired-end 150-bp reads. Adapters and low-quality bases (Phred < 20) were removed from the paired-end 150-bp reads with Trim Galore v0.6.10 (https://github.com/FelixKrueger/TrimGalore; doi:10.5281/zenodo.7598955), a wrapper around Cutadapt (Martin, 2011). We used default settings: automatic adapter detection, and read pairs discarded if either read was shorter than 20 bp. Trimmed reads were aligned with bowtie2 v2.5.5 (Langmead & Salzberg, 2012) using ‘--very-sensitive -X 2000 --no-mixed --no-discordant’ to ValleyOak3.2 plus the v3.0 chloroplast and mitochondrial contigs. Overall alignment rates were 46.8% (786_D1) and 97.1% (786_D3). Duplicates were marked with samtools v1.24 (Danecek *et al*., 2021) using ‘markdup’ and excluded, and we kept properly paired reads on nuclear sequences with MAPQ ≥ 30. Peaks were called with MACS3 v3.0.4 (Doherty *et al*., 2026) using ‘callpeak -f BAMPE --keep-dup all -q 0.01’ with an effective genome size of 844,024,183 bp for each library and for the pooled reads. Reproducible peaks were pooled peaks that overlapped a peak in each library by ≥ 50% of either peak using bedtools v2.31.1 (Quinlan & Hall, 2010), after removing the chromosome 1 mitochondrial mis-assembly. A genomic bin was classified as accessible when located within 500 bp of a reproducible peak, with 5,113 total peaks (median 466 bp).

### Oxford Nanopore PromethION genome and methylome sequencing

High molecular weight genomic DNA was extracted from half of each young leaf sample following the cetyltrimethylammonium bromide (CTAB) method described by (Inglis *et al*., 2018), with the following modifications: (i) sodium metabisulfite (1% w/v) was used instead of 2-mercaptoethanol (1% v/v) in the sorbitol wash buffer and CTAB solution; (ii) tissue homogenate wash steps were repeated until the supernatant became clear; (iii) the CTAB lysis step was performed at 45°C; and (iv) the chloroform extraction step was performed twice using ice-cold chloroform. DNA purity was assessed by absorbance ratios measured using the NanoDrop ND-1000 spectrophotometer (Thermo Fisher Scientific, MA). DNA yield was quantified using the QuantiFluor ONE dsDNA Dye assay (Promega, WI). The size distribution of the high molecular weight DNA was estimated using the Femto Pulse system with the Genomic DNA 165 kb kit (Agilent, CA).

Libraries were sequenced at University of California Davis DNA Technologies core facility (Davis, CA) using a PromethION flow cell for 48 h. We achieved between 11.8 - 28.1x genome coverage per sample (Figure S1C). Nucleotide bases and all-context methylation were called from the raw sequence data using an A5000 GPU box with Dorado 0.5.0, after which sequencing data was aligned to the valley oak reference genome (accession GCF_001633185.2), retaining modified-base calls in MM/ML tags. Cytosine methylation was quantified using modkit v0.6.4 ‘pileup’ separately for CG, CHG, and CHH contexts. We retained primary alignments with mapping quality ≥10 and combined 5mC and 5hmC calls. A common modification-confidence threshold (0.8027), determined from the median of per-sample ‘modkit sample-probs’ estimates, was applied to all samples. CG calls were combined across strands, whereas CHG and CHH calls were retained by strand. Analyses were restricted to the 12 assembled nuclear chromosomes (812.7 Mb). The raw sequence data has been deposited in the International Nucleotide Sequence Database Collaboration (INSDC) database under the BioProject accession number XXXXXXX.

Gene models were obtained from the ValleyOak3.2 RefSeq annotation. Genomic regions were classified as gene body, strand-aware upstream (≤1.5 kb), downstream (≤1.5 kb), or intergenic, using that order of precedence for overlapping annotations. We obtained TE annotations from the same reference genome (Sork *et al*., 2022), using the sequence identity clusters (45% similarity; s1RF* families) with copy number defined as the number of annotated copies in each cluster. Where a bin overlapped multiple TEs, the TE with the greatest overlap was designated its primary TE.

### Drought-enhanced methylation regions

#### Identification of drought-enhanced methylation regions

We followed established approaches to identifying drought-enhanced methylation regions (Zhong *et al*., 2021; Miryeganeh *et al*., 2022). We identified methylation changes in fixed, non-overlapping 200-bp genomic bins. Cytosines were included when covered by ≥3 reads, and bins were retained when they contained ≥32 covered CHH cytosines to compensate for low methylation levels in CHH, or ≥4 covered CG or CHG cytosines in every sample, as in (Zhang *et al*., 2026). These criteria yielded 2,692,613 CHH, 2,130,162 CHG, and 798,463 CG bins.

For each treatment and interval (T1-T2, T1-T3 and T2-T3), we identified regions showing consistent methylation changes that exceeded those in the opposite treatment. A bin was required to meet four criteria: (1) Group-level change: A significant change in methylation counts pooled across the four plants (Fisher’s exact test, Benjamini-Hochberg FDR <0.05), with an absolute methylation difference of ≥0.10 for CHH, ≥0.20 for CHG or ≥0.40 for CG. (2) Consistency across plants: A change in the same direction in all four plants, each exceeding half the corresponding group-level effect-size threshold. (3) Treatment × time interaction: A significant difference between treatments in the pooled methylation odds ratios across time points (Woolf’s test with a Haldane correction, FDR <0.05), in the direction of the focal treatment. (4) Greater response than the opposite treatment: A difference in mean per-plant methylation change between treatments of ≥0.05 for CHH, ≥0.10 for CHG or ≥0.20 for CG, in the focal treatment’s direction. Adjacent qualifying bins with the same direction of change were merged into regions. Importantly, regions changing in both treatments were retained when the focal treatment exhibited a greater response. Applying the same criteria with treatments reversed identified control-enhanced regions. We interpreted region counts in relation to the reciprocal control-enhanced set and performed an independent plant-blocked DSS analysis to assess the robustness of the principal findings (Supplementary Methods).

#### Timing and consistency of methylation changes

To characterise the timing of CHH gains, we followed drought-enhanced T1-T3 regions across all three time points, calculating methylation changes from pooled calls within each region and averaging first across regions within plants, then across plants within treatments. The relative contribution of T2-T3 was calculated from the drought-control difference in methylation change during each interval. T2 measurements were not used to select these regions.

To assess consistency independently of region selection, we repeated DMR calling four times, each time excluding one drought plant from all selection steps and requiring concordant changes in the remaining three. We then measured the excluded plant’s T1-T3 methylation change at the resulting regions and compared it with changes at the full DMR set. To assess the spatial extent of methylation gains, we also examined tested bins within 1 kb of, but outside, the identified regions.

### Genomic characteristics of DNA methylation gain

We investigated whether drought-enhanced CHH gains preferentially occurred in particular genomic contexts while accounting for differences in the probability of detecting methylation change. Because baseline methylation and sequencing depth influence both the measurable magnitude and statistical power of methylation changes, all enrichment analyses used tested 200-bp bins as the background. We first grouped bins into detection strata defined by deciles of T1 CHH methylation and quintiles of sequencing depth. For each genomic feature, observed overlap with drought-enhanced CHH-gain bins was compared with 1,000 random sets of tested bins matched across these strata. Fold enrichment was calculated as observed divided by mean expected overlap, with empirical *P* values adjusted using the Benjamini-Hochberg procedure within feature categories.

To identify genomic properties independently associated with CHH gain, we fitted multivariable logistic regressions to tested TE-overlapping bins. The outcome was whether a bin overlapped a drought-enhanced CHH-gain region. Predictors included detection stratum, TE class, log10 TE length, log10 family copy number, log10 distance to the nearest gene, genic context and proximity to an accessible chromatin region (identified at T1, ≤500 bp). Continuous predictors were standardised and effect sizes reported as odds ratios per standard deviation. Models included all gain bins and a random sample of non-gain bins. We additionally estimated TE enrichment using stratified odds ratios and alternative adjustments for baseline methylation and depth. Sensitivity analyses used one randomly selected bin per region to reduce non-independence between adjacent bins. Because multiple bins may belong to the same TE or region, conventional Wald confidence intervals may underestimate uncertainty; we therefore emphasised effect sizes and consistency across analyses. Equivalent models were fitted to control-enhanced CHH gains.

### Shared methylation targets between treatments

To determine whether individual plants gained CHH methylation at the same TE copies more often than expected from their genomic characteristics, we identified gains independently within each treatment. A bin was classified as gaining methylation if it showed a significant pooled increase (Fisher’s exact test, FDR < 0.05; ΔCHH ≥ 0.1) and an increase of ≥ 0.05 in all four plants. We excluded the between-treatment criteria used to identify drought-enhanced regions, as these would restrict overlap between treatments by design. Analyses were performed on tested CHH bins overlapping annotated TEs and repeated at the level of individual TE copies, classified as gaining methylation if any constituent bin gained.

We tested whether control gains predicted drought gains between T1-T3 using logistic regression, adjusting for baseline methylation, sequencing depth, TE class, length, copy number, gene proximity, genic context, and accessible chromatin regions. We also compared the observed overlap with that expected from these genomic characteristics alone and examined whether control plants showed greater methylation changes at drought-gaining bins than at matched non-gaining bins.

### RNA extraction, sequencing, and orthology assignment

Six seedlings had transcriptomes at both T1 and T3 (four drought-treated and two well-watered), of which three were also included in the methylome dataset. Leaf tissue was frozen in liquid nitrogen and ground with beads. To extract RNA, we added 1.8 mL cold extraction buffer to each tube consisting of 8 M urea, 3 M LiCl, 1.76% polyvinylpyrrolidone K-60 solution, and 10 mM dithiothreitol (added from 1 M stock just before use). Tubes were vortexed vigorously and ground for 10 secs at 30 Hz, followed by centrifugation for 10 min at 1,000 rcf and 4° C that allowed recovery of 1.4 mL supernatant, which was kept at 4° C for twenty minutes. Next, we centrifuged tubes for 30 min at 20,000 rcf and 4 °C. After discarding the supernatant, the pellet was washed with 2 rounds of 70% ethanol, each followed by a 5-min centrifugation at 5,000 rcf and room temperature. Finally, the pellet was dried in the fume hood for 10 min. The air-dried pellet was used as the starting material for extraction of total RNA with the RNeasy Plant Mini Kit (QIAGEN, Germantown, MD). RNA was quantified using a NanoDrop 2000 (Thermo Fisher), with the RNA quality assessment (Qubit and Fragment Analyzer), library preparation, and Next Generation Sequencing performed by DNA Technologies and Expression Analysis Core at the UC Davis Genome Center.

Libraries were prepared with 3′ Tag-Seq and sequenced as single-end 101-bp reads, 8-10 million reads per library. Because 3′ Tag-Seq relies on oligo-dT priming, expression estimates reflect polyadenylated transcripts. Reads were quality- and adapter-trimmed with Trimmomatic v0.39 (ILLUMINACLIP:TruSeq3-PE.fa:2:30:10 SLIDINGWINDOW:4:5 LEADING:5 TRAILING:5 MINLEN:25) (Bolger *et al*., 2014) and checked with FastQC v0.11.2 (Andrews, 2010). Ribosomal RNA reads were removed by mapping to the SILVA rRNA database (Quast *et al*., 2012) with BBMap (Bushnell, 2014) and keeping only unmapped reads with ‘outu.’ Reads were aligned to ValleyOak3.2 (GCF_001633185.2) with STAR v2.7.11b (Dobin *et al*., 2013) using default settings, except that we kept reads mapping to up to 100 loci ‘-- outFilterMultimapNmax 100’. All alignments of multi-mapping reads were retained, so each contributed coverage at every locus it mapped to; this avoids systematically excluding transcripts from repetitive regions. Unique molecular identifiers were not used, so PCR duplicates were not removed. Coverage was normalised to counts per million (CPM) in 50-bp bins with deepTools bamCoverage with ‘--normalizeUsing CPM’ (Ramírez *et al*., 2016). Gene-level expression was the sum of coverage values over each annotated gene body on the 12 chromosomes, computed with bedtools map ‘-o sum’ (Quinlan & Hall, 2010). Expression was analysed as log2(CPM + 1).

For functional analyses, each oak gene was assigned to the top SwissProt BLASTP hit from the Trinotate annotation (Bryant *et al*., 2017), keeping the lowest-e-value hit across a gene’s transcripts. Genes whose top hit was an *Arabidopsis thaliana* protein were mapped to TAIR loci through the UniProt ID mapping (UniProt Consortium, 2023). Gene ontology (GO) terms were taken from org.At.tair.db (Carlson, 2026), and oak paralogues mapping to the same TAIR locus were collapsed for GO tests.

### Gene expression near CHH-gaining transposable elements

Genes were retained when expressed (CPM > 0) at both time points in all six plants. For each gene, we calculated the mean T1-T3 log2 expression change in drought plants minus the corresponding mean change in well-watered controls, termed the interaction effect (IE). Genes were classified as induced (IE ≥1) or repressed (IE ≤−1). Because of the small and unbalanced number of biological replicates (four drought-treated and two well-watered plants), we classified expression responses using an effect-size threshold rather than statistical significance testing. These categories describe estimated drought-associated expression responses rather than statistically established differential expression.

A gene was associated with a CHH-gaining TE when a drought-enhanced T1-T3 CHH region overlapped an annotated TE within 1.5 kb upstream or 1.5 kb downstream. Analyses were restricted to genes with at least one tested CHH bin within a TE in that window. Logistic regression tested whether TE gain was associated with induction or repression, adjusting for gene length, baseline expression, and the number of tested TE bins. We additionally tested whether ABA-response and water-deprivation genes were more likely to carry CHH-gaining TEs than other induced genes. Restricting these analyses to induced genes accounted for the general enrichment of stress-response functions among drought-induced genes. Models adjusted for gene length, baseline expression, interaction effect, the number of tested TE bins, and baseline CHH methylation within the window. ABA-response and water-deprivation genes were identified through Arabidopsis orthologues annotated to GO:0009737 and GO:0009414, respectively. Analyses were repeated after collapsing oak paralogues to Arabidopsis orthologues.

### Candidate new TE insertions

We searched for candidate somatic TE insertions in ONT reads using a custom read-level pipeline. Insertions ≥50 bp were identified from CIGAR insertions or split-read alignments, requiring ≥1 kb of high-quality (MAPQ 60) alignment on both flanks. Inherited insertions were excluded using a catalogue generated by Sniffles2 (Smolka *et al*., 2024) across all 24 samples. Candidate inserted sequences were classified using RepeatMasker, screened against organelle sequences and clustered across samples by genomic position and insertion length. Candidates supported by multiple independent molecules were re-genotyped across all samples to identify events present in individual leaves but absent from the same plant’s other sampled leaves. Non-TE insertions detected using the same procedure provided an internal estimate of library artefacts. Insertion rates were normalised by callable sequence per library and compared between treatments using each plant’s change from T1 to later leaves. Treatment differences were assessed using an exact permutation test over the 70 possible assignments of eight plants into two groups of four.

## Data analysis

The following R packages were used: R 4.6.1 (R Core Team): DSS 2.60.0 (Park & Wu, 2016), bsseq 1.48.0 (Hansen *et al*., 2012), data.table 1.18.6.1 (Barrett *et al*., 2024), tidyverse 2.0.0 (Wickham *et al*., 2019), GenomicRanges 1.64.0 (Lawrence *et al*., 2013), GenomeInfoDb 1.40.1 (Lawrence *et al*., 2013), Biostrings 2.80.2 (Pages *et al*., 2024a), Rsamtools 2.28.0 (Morgan *et al*., 2024), AnnotationDbi 1.66.0 (Pages *et al*., 2024b), org.At.tair.db 3.19.1 (Carlson, 2026), GO.db 3.19.1 (Carlson, 2024), cowplot 1.1.3 (Wilke, 2024).

## Data availability

Raw Oxford Nanopore reads (with basecalled modification tags), 3′ Tag-Seq reads and ATAC-seq reads have been deposited in the NCBI Sequence Read Archive under BioProject PRJNAXXXXXX. Processed data are available in Zenodo (DOI), including methylation calls (bedMethyl files), differentially methylated region sets, gene-level expression tables, ATAC-seq peaks and physiological measurements. Source data for all figures are provided with this paper and in the same repository. The *Q. lobata* reference genome (ValleyOak3.2, GCF_001633185.2) is available from NCBI.

## Code availability

All custom analysis and plotting scripts, and the bioinformatic workflows, are available in Zenodo (DOI) and on GitHub (URL).

## Author contributions

LDP and VLS planned and designed the research. LDP performed experiments and analysed data. LDP and VLS wrote the manuscript.

## Acknowledgements

This work was performed (in part) at UC Nature’s UCSB Sedgwick Reserve, doi:10.21973/N3C08R. We acknowledge the Chumash People as the traditional caretakers (past, present, and future) of the land where this research was conducted. The sequencing was carried out by the DNA Technologies and Expression Analysis Core at the UC Davis Genome Center, supported by NIH Shared Instrumentation Grant 1S10OD010786-01. The analyses used computational and storage services associated with the Hoffman2 Cluster, which is operated by the UCLA Office of Advanced Research Computing’s Research Technology Group. Thanks to A. Lentz for collecting the acorns and S. Cokus for valuable discussions about transposable element biology. For financial support, we thank The Seaver Institute and the National Science Foundation (IOS-2512127).

