## Supplementary material for "Repeated drought amplifies DNA methylation at shared transposable element targets in *Quercus lobata*": NP-MS-supp.pdf

L. D. Peck* and V. L. Sork*

This PDF file includes:
Supplementary Methods
Supplementary Results
Figs. S1 to S5
Tables S1 to S6

### Supplementary Methods

#### Plant-blocked DSS analysis

As a complementary analysis accounting for biological replication, we fitted DSS 2.60.0 (Wu *et al.*, 2013, 2015; Feng *et al.*, 2014; Park & Wu, 2016) to the same 200-bp bins and cytosine coverage filters used in the primary analysis. Methylated and total cytosine calls were summed within each bin separately for each sample. All 24 samples were analysed jointly using 'DMLfit.multiFactor' with the design ~ plant + time + drought × T2 + drought × T3'. Plant was an eight-level factor accounting for individual baseline methylation differences, including those associated with maternal family and treatment assignment, and time was a three-level factor. We adjusted *P* values across all tested bins separately for each contrast using the Benjamini-Hochberg procedure. Adjacent significant bins (FDR < 0.05) with effects in the same direction were merged into regions. Regional methylation effect sizes were calculated from raw counts as the difference between mean methylation changes in drought-treated and well-watered plants. Individual-plant changes were also calculated. CG and CHG methylation were analysed using the same approach. To assess the sensitivity of regional CHH responses to genome-wide methylation shifts, we refitted the model including each sample's genome-wide mean CHH methylation (arcsine square-root transformed and centred) as an additional covariate. Regions identified by the two models were compared by genomic overlap. Because genome-wide methylation may itself respond to drought, this analysis assesses sensitivity to global methylation variation rather than treating such variation solely as a confounding factor.

DSS regions were annotated and their genomic characteristics compared with the background of tested bins using the same detection-matched resampling procedure as the primary analysis (T1 methylation decile × sequencing-depth quintile; 1,000 resamples; Benjamini-Hochberg correction within feature categories). Baseline chromatin accessibility of TEs containing DSS regions was compared with that of matched TEs using continuous Tn5 cut density. Genes with a DSS region within 1.5 kb upstream were compared with other genes containing tested CHH bins in this window, accounting for the number of tested bins and their baseline methylation. Drought-associated expression changes were quantified as the difference between treatment groups in T1-T3 expression changes using all available T1 and T3 libraries (nine plants). A pre-treatment comparison of expression between treatment groups served as a negative control.

Finally, DSS regions were intersected with regions identified using the primary region-calling approach. For non-overlapping DSS regions, we examined pooled methylation changes in drought-treated plants to determine whether these regions fell below the primary CHH effect-size threshold of 0.10.

### Supplementary Results

#### Drought-induced DNA methylation using a plant-blocked DSS analysis

To assess whether our findings were sensitive to the region-calling approach, we fitted a plant-blocked DSS model (methylation ~ plant + time + drought × T2 + drought × T3) to 200-bp bins. We analysed 2,692,613 CHH bins containing ≥ 32 cytosines covered by ≥ 3 reads in all 24 samples, testing drought × time interactions at FDR < 0.05. Blocking on plant accounted for individual baseline methylation differences, allowing us to identify regions with differential temporal responses between treatments.

The DSS analysis reproduced the delayed CHH response observed in the primary analysis (Table S1). Only one region was significant between T1 and T2 (a loss), compared with 188 between T1 and T3 (183 gains, five losses) and 93 between T2 and T3 (92 gains). At 97% of T1-T3 regions, all four drought-treated plants gained methylation, with mean gains of 7.0-8.7 percentage points across individuals, including the weakest genome-wide responder (786_D2). Well-watered plants showed comparatively little change (−1.9 to +0.4 percentage points). Including genome-wide mean CHH methylation as a covariate retained 122 T1-T3 regions (107 gains) and 94 T2-T3 regions (84 gains), of which 81 T1-T3 regions overlapped those identified by the original DSS model. Therefore, the response was not explained solely by changes in genome-wide methylation levels.

The genomic distribution of DSS regions broadly supported the primary analysis. After matching, T1-T3 regions overlapped TEs (Mantel-Haenszel odds ratio = 1.44), which is in the same direction as the primary analysis, but weaker (odds ratio = 2.5-3). The T2-T3 regions showed no TE excess (odds ratio = 0.95). In addition, CHH gains were enriched upstream of genes, particularly between T2 and T3 (2.5-fold, FDR = 0.014; 2.9-fold after adjustment for genome-wide methylation, FDR = 0.007). This enrichment was predominantly attributable to upstream TEs (2.3-2.6 fold, FDR ≤ 0.02). T1-T3 gains were also enriched upstream (1.6 fold; 2.1 fold after adjustment, FDR = 0.028), with additional downstream enrichment in some models (T2-T3: 2.3-fold, FDR = 0.039). Intergenic regions were depleted (0.71-0.83 fold), while gene bodies showed no enrichment. TEs containing DSS regions also had greater T1 chromatin accessibility than matched TEs (mean log2 accessibility 0.55 vs 0.42, $P$ = 0.028; 0.69 vs 0.48 after genome-wide adjustment, $P$ = 0.008).

Other methylation contexts showed comparatively few significant regions (CG: 0-9; CHG: 29 at T1-T3, approximately equally divided between gains and losses). Genes associated with DSS regions showed no detectable drought-related expression changes. Finally, the two approaches identified largely different regions: 39 of the 188 T1-T3 DSS regions overlapped those identified by the primary 4/4 pattern-based approach. Most of the remainder fell below the primary analysis' 10-pp pooled change threshold (median DSS change = 7.7 pp). These analyses support the principal findings of a delayed, predominantly CHH-specific methylation gain, concentrated near genes and in initially accessible chromatin, that is robust to alternative region-calling methods and adjustment for genome-wide methylation.

A B

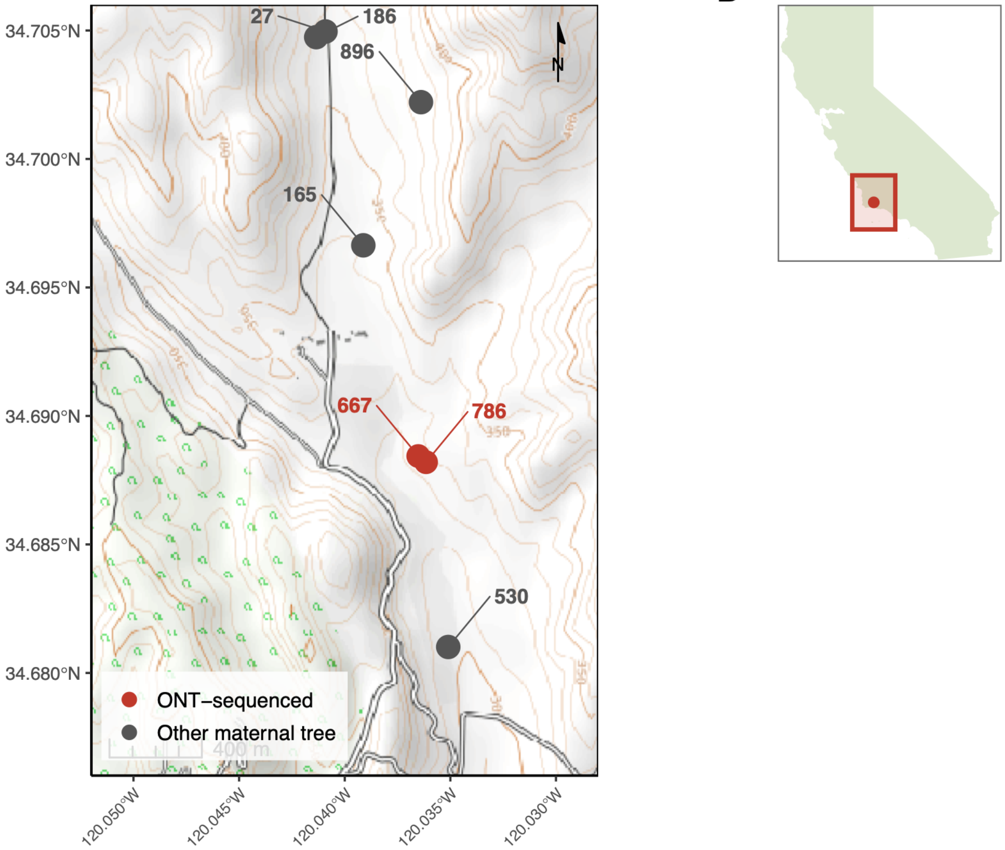


C

| Individual | Family | Treatment | Mean depth T1 | Mean depth T2 | Mean depth T3 | Mean background CG (M) | Mean background CHG (M) | Mean background CHH (M) |
|---|---|---|---|---|---|---|---|---|
| 667_C1 | 667 | Control | 16.4 | 16.4 | 23.3 | 21.8 | 31.0 | 207.1 |
| 667_C2 | 667 | Control | 20.6 | 22.3 | 19.3 | 21.8 | 30.9 | 206.7 |
| 667_D1 | 667 | Drought | 11.8 | 28.1 | 14.3 | 21.6 | 30.6 | 205.1 |
| 667_D2 | 667 | Drought | 17.3 | 13.3 | 17.7 | 21.7 | 30.8 | 206.4 |
| 786_C1 | 786 | Control | 23.6 | 19.1 | 20.4 | 23.3 | 32.7 | 217.1 |
| 786_C3 | 786 | Control | 25.5 | 18.6 | 14.9 | 23.3 | 32.7 | 217.5 |
| 786_D2 | 786 | Drought | 22.7 | 19.3 | 15.6 | 23.2 | 32.5 | 216.2 |
| 786_D3 | 786 | Drought | 16.00 | 14.3 | 22.5 | 23.2 | 32.5 | 216.1 |

**Figure S1. Study site and sequenced individuals.** (A) Map of the locations of seven *Quercus lobata* trees sampled for acorn collection at the UC Santa Barbara Sedgwick Reserve, Santa Barbara County, California. Trees from two maternal families (667 and 786; red circles) were selected for Oxford Nanopore sequencing; remaining maternal trees (black circles) contributed seedlings to the drought phenotyping experiment only. Basemap: OpenTopoMap. (B) Location of Sedgwick Reserve  within map of California. (C) Oxford Nanopore sequencing metrics for eight seedlings collected from the 667 and 786 maternal trees and which were included in methylation analysis (four per maternal family × two treatments). Mean depth represents the average read depth across cytosine sites for the CG context (CHG and CHH values are similar). Mean background represents site counts averaged across T1-T3 comparisons. M = million cytosine sites.

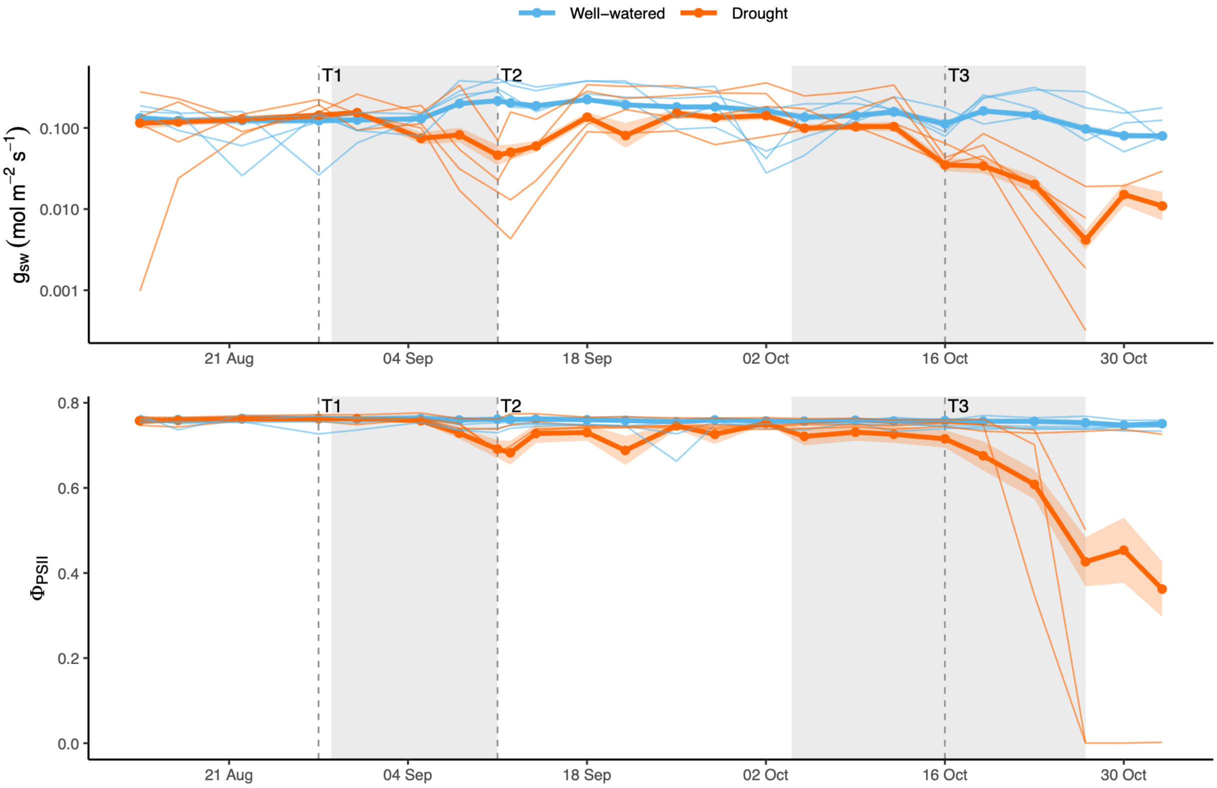


**Figure S2. Drought treatment reduced stomatal conductance and, late in the second drought, photosystem II efficiency.** (A) Stomatal conductance (gsw, log scale) and (B) operating efficiency of photosystem II (ΦPSII) of all seedlings in the drought experiment (80 seedlings from seven maternal trees; 40 drought, 40 well-watered), measured with a LI-600 porometer. Thick lines and ribbons: treatment mean ± SE across seedlings (gsw: geometric mean, SE on the log scale); thin lines: the eight ONT-sequenced seedlings. Grey shading, water withheld from the drought group (30 Aug-11 Sep and 4-27 Oct 2023); dashed lines, leaf sampling for ONT sequencing (T1, T2, T3). ΦPSII fell only after T3, so T3 leaves were sampled under strong stomatal limitation but before loss of photochemical efficiency.

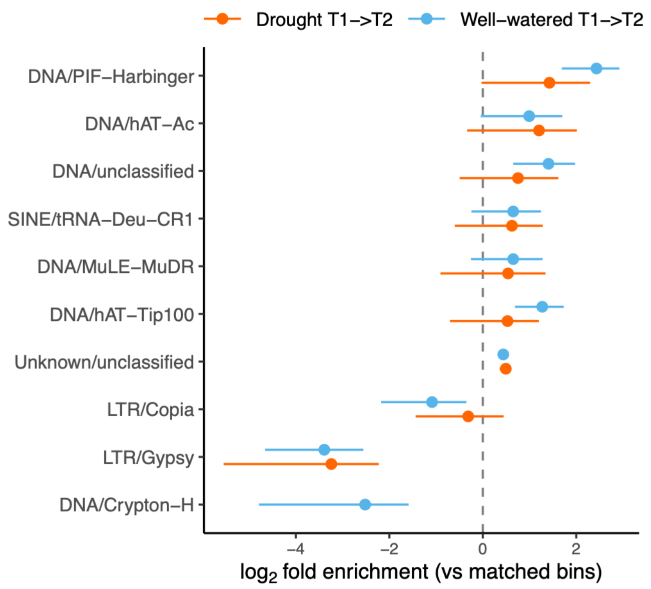


**Figure S3. Well-watered CHH gains favour the same TE classes as drought gains at T2.** Fold enrichment (log2) of CHH gain bins in each TE superfamily relative to bins matched on baseline (T1) CHH methylation and read depth, for T1→T2 gain regions in drought (orange) and well-watered (blue) plants. Points, fold enrichment; bars, 95% CIs from a 1 Mb genomic block bootstrap. Dashed line, no enrichment.

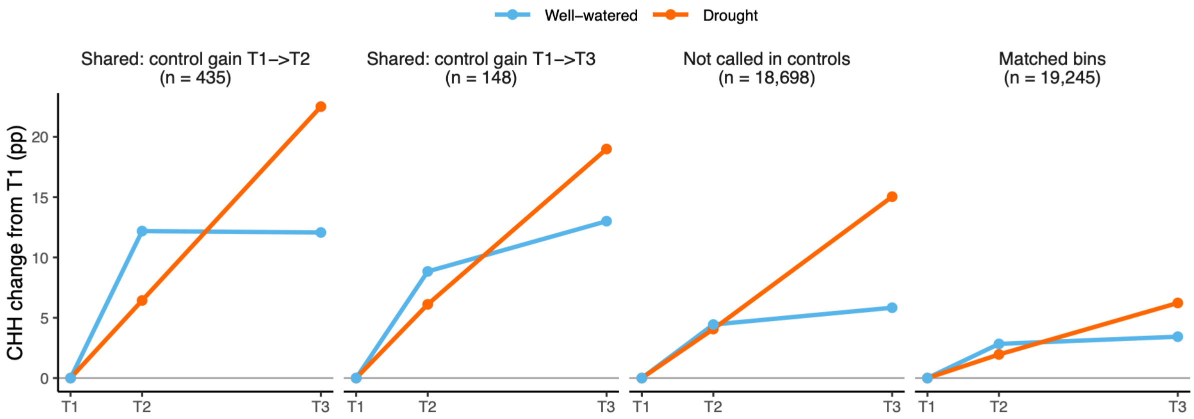


**Figure S4. At shared targets, well-watered plants gain early and plateau while drought plants keep gaining.** Mean CHH change from T1 (percentage points) in drought (orange) and well-watered (blue) plants at drought T1→T3 gain bins (200-bp bins that are a pooled-read Fisher DMR in the drought group, BH-adjusted, |Δ| ≥ 0.1, with all four drought plants gaining ≥ 5 pp. These bins were selected before the interaction test and difference-in-differences filters, which require drought gain to exceed control gain and so would bias the overlap with control gain downwards. Bins were split by whether well-watered plants also gained there (T1→T3 or T1→T2) or not, and at bins that were matched by T1 methylation and sequencing depth.

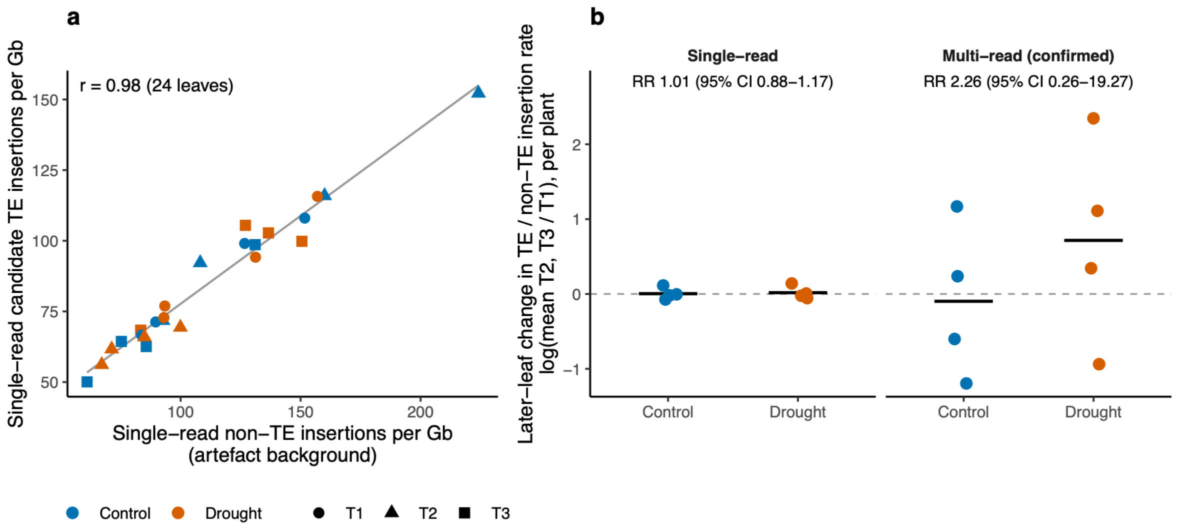


**Figure S5. No detectable increase in candidate new TE insertions under drought**. (a) Per-leaf rates of single-read candidate TE insertions and non-TE insertions, the latter providing an estimate of library-specific technical background (24 leaves; colour indicates treatment, shape indicates timepoint; line shows the least-squares fit). (b) Plant-level changes in TE insertion rates, normalised by the corresponding non-TE insertion rates, between the pre-treatment leaf (T1) and leaves formed during the experiment (mean of T2 and T3). Results are shown for single-read candidates and multi-read candidates retained after cross-leaf re-genotyping. Points represent individual plants (n = 4 per treatment); horizontal bars indicate treatment means and the dashed line indicates no temporal change. RR is the ratio of temporal changes between drought and well-watered plants, with 95% confidence intervals calculated using Welch's method on log-transformed changes and back-transformed to the rate-ratio scale.

**Table S1. Number of CHH, CHG and CG regions gaining or losing methylation in each interval.** Drought and well-watered columns: 4/4 pattern rule applied to each group in turn (pooled-read Fisher test, BH-adjusted, $|\Delta| \geq 0.1$; all four plants changing ≥ 5 pp in the same direction; read-level treatment × time interaction; plant-mean difference-in-differences ≥ 5 pp relative to the other group). CHH thresholds; CHG and CG thresholds are 2× and 4×. DSS columns: plant-blocked DSS treatment × time test on 200-bp bins, FDR < 0.05 (gain = higher in drought).

| Context | Interval | Drought gain | Drought loss | Well-watered gain | Well-watered loss | DSS DMRs gain | DSS DMRs loss |
|---|---|---|---|---|---|---|---|
| CHH | T1→T2 | 264 | 0 | 404 | 0 | 0 | 1 |
| CHH | T1→T3 | 14055 | 3 | 22 | 4 | 183 | 5 |
| CHH | T2→T3 | 5835 | 5 | 2 | 6 | 92 | 1 |
| CHG | T1→T2 | 14 | 29 | 3 | 8 | 1 | 2 |
| CHG | T1→T3 | 42 | 315 | 7 | 16 | 14 | 15 |
| CHG | T2→T3 | 29 | 117 | 17 | 8 | 6 | 11 |
| CG | T1→T2 | 0 | 0 | 0 | 0 | 0 | 0 |
| CG | T1→T3 | 0 | 9 | 0 | 0 | 0 | 0 |
| CG | T2→T3 | 0 | 2 | 0 | 0 | 1 | 8 |

**Table S2. Overlap of drought and well-watered CHH-gain TE bins (200 bp).** Gain bins: pooled-read Fisher DMR of that group (BH, |Δ| ≥ 0.1) with all four plants gaining ≥ 5 pp, unless stated. Expected: overlap predicted from T1 CHH × depth strata, TE class, length, copy number, genic context and ATAC. Observed / expected 95% intervals (1 Mb genomic block bootstrap) are in Supplementary Data 3. The pattern rule requires drought gain to exceed control gain and is therefore biased towards less overlap. Low-copy: TE families at or below the median copy number (≤ 2,118 copies), a mappability proxy.

| Comparison | Drought-gain bins | Well-watered-gain bins | Shared, observed | Shared, expected | Observed / expected |
|---|---|---|---|---|---|
| Drought T1→T3 vs well-watered T1→T3 | 19255 | 588 | 148 | 65 | 2.29 |
| Drought T1→T3 vs well-watered T1→T2 | 19255 | 1345 | 435 | 171 | 2.54 |
| Drought T1→T3 vs well-watered T1→T3, well-watered pooled-read rule only | 19255 | 6146 | 2152 | 765 | 2.81 |
| Drought T1→T3 pattern rule vs well-watered T1→T3 | 15208 | 588 | 72 | 51 | 1.4 |
| Drought T1→T3 vs well-watered T1→T3, low-copy TE families | 5622 | 140 | 34 | 15 | 2.33 |
| Drought T1→T3 vs well-watered T1→T2, low-copy TE families | 5622 | 362 | 116 | 44 | 2.65 |

**Table S3. Tests of CHH gain in expressed genes.** We tested CHH gain in the 1.5 kb upstream of genes (TE-overlapping drought T1→T3 gain regions) against drought expression change (induced / repressed: Interaction effect rule) and against ABA-response and water-deprivation GO terms among induced genes.

| Question | Test | Genes (n) | OR | *P* |
|---|---|---|---|---|
| Induced in drought ~ upstream CHH gain | All tested genes (Fisher) | 22169 | 1.09 | 0.272 |
| Induced in drought ~ upstream CHH gain | Genes with an upstream TE (Fisher) | 18034 | 1.05 | 0.568 |
| Induced in drought ~ upstream CHH gain | Genes with an upstream TE, adjusted for gene length, T1 expression and n upstream TE bins | 18034 | 1.22 | **0.018** |
| Repressed in drought ~ upstream CHH gain | All tested genes (Fisher) | 22169 | 0.87 | 0.087 |
| Repressed in drought ~ upstream CHH gain | Genes with an upstream TE (Fisher) | 18034 | 0.90 | 0.243 |
| Repressed in drought ~ upstream CHH gain | Genes with an upstream TE, adjusted for gene length, T1 expression and n upstream TE bins | 18034 | 0.98 | 0.801 |
| GO: response to abscisic acid (GO:0009737) | Induced genes: term ~ upstream gain (Fisher) | 788 | 1.88 | 0.074 |
| GO: response to abscisic acid (GO:0009737) | Induced genes with an upstream TE (Fisher) | 698 | 1.70 | 0.124 |
| GO: response to abscisic acid (GO:0009737) | Induced genes with an upstream TE: term ~ gain, adjusted (length, T1 expression, induction, n upstream TE bins) | 698 | 1.77 | 0.120 |
| GO: response to abscisic acid (GO:0009737) | Genes with an upstream TE: induced × gain interaction | 10038 | 1.66 | 0.169 |
| GO: response to abscisic acid (GO:0009737) | Induced genes with an upstream TE: gain ~ term + detection covariates (primary) | 698 | 1.74 | 0.143 |
| GO: response to water deprivation (GO:0009414) | Induced genes: term ~ upstream gain (Fisher) | 788 | 1.31 | 0.487 |
| GO: response to water deprivation (GO:0009414) | Induced genes with an upstream TE (Fisher) | 698 | 1.21 | 0.647 |
| GO: response to water deprivation (GO:0009414) | Induced genes with an upstream TE: term ~ gain, adjusted (length, T1 expression, induction, n upstream TE bins) | 698 | 1.15 | 0.764 |
| GO: response to water deprivation (GO:0009414) | Genes with an upstream TE: induced × gain interaction | 10038 | 1.12 | 0.802 |
| GO: response to water deprivation (GO:0009414) | Induced genes with an upstream TE: gain ~ term + detection covariates (primary) | 698 | 1.21 | 0.691 |

**Table S4. ABA-response and water-deprivation genes induced in drought plants with a TE-overlapping CHH-gain region within 1.5 kb upstream.** CHH gain: largest drought – control change among the gene's upstream regions. Expression change: mean drought log2 fold change minus mean well-watered log2 fold change (T1→T3).

| Oak gene | Arabidopsis orthologue | Symbol | GO term | Upstream TE superfamily | Gain interval | CHH gain (drought-control, pp) | Expression change (drought-control, log2) |
|---|---|---|---|---|---|---|---|
| LOC115964092 | AT1G20440 | AtCOR47/COR47/RD17 | ABA response; water deprivation | DNA/MuLE-MuDR | T1→T3 | 9.9 | 4.42 |
| LOC115976285 | AT3G29575 | AFP3 | ABA response | Unknown/unclassified | T1→T3 | 11.9 | 3 |
| LOC115977240 | AT4G30960 | ATCIPK6/CIPK6/SIP3/SNRK3.14 | water deprivation | Unknown/unclassified | T1→T3 | 20.1 | 2.25 |
| LOC115967588 | AT2G22570 | ATNIC1/NIC1 | ABA response | Unknown/unclassified | T1→T3 | 18.6 | 1.79 |
| LOC115982679 | AT5G13330 | Rap2.6L | ABA response; water deprivation | Unknown/unclassified | T1→T3 | 14.5 | 1.74 |
| LOC115974175 | AT1G45249 | ABF2/AREB1/AtABF2/ATAREB1 | ABA response; water deprivation | Unknown/unclassified | T1→T3 | 6.5 | 1.65 |
| LOC115967252 | AT5G42900 | COR27 | ABA response | | T1→T3 | 6.9 | 1.53 |
| LOC115952703 | AT2G35940 | BLH1/EDA29 | ABA response | DNA/MuLE-MuDR | T1→T3 | 14.4 | 1.51 |
| LOC115970777 | AT5G05410 | DREB2/DREB2A | water deprivation | DNA/unclassified | T1→T3 | 10.6 | 1.48 |
| LOC115993038 | AT5G01520 | AIRP2/AtAIRP2 | ABA response | Unknown/unclassified | T1→T3 | 21 | 1.41 |
| LOC115969855 | AT1G74120 | mTERF15 | ABA response | DNA/hAT-Ac | T1→T3 | 9.1 | 1.41 |
| LOC115992535 | AT1G29395 | COR413-TM1/COR413IM1/COR414-TM1/DEG25 | ABA response; water deprivation | DNA/hAT-Tip100 | T2→T3 | 9.2 | 1.33 |
| LOC115974673 | AT2G46270 | GBF3 | ABA response | Unknown/unclassified | T1→T3 | 16.2 | 1.22 |
| LOC115961008 | AT5G46050 | AtNPF5.2/ATPTR3/NPF5.2/PTR3 | ABA response | DNA/unclassified | T1→T3 | 7.7 | 1.14 |
| LOC115974684 | AT4G35300 | TMT2 | water deprivation | Unknown/unclassified | T1→T3 | 19.1 | 1.13 |
| LOC115965831 | AT1G74120 | mTERF15 | ABA response | Unknown/unclassified | T1→T3 | 7.6 | 1.12 |
| LOC115970748 | AT3G22380 | TIC | ABA response; water deprivation | Unknown/unclassified | T1→T3 | 11.2 | 1.09 |
| LOC115977558 | AT3G13460 | ECT2 | ABA response | Unknown/unclassified | T1→T3 | 8.2 | 0.92 |
| LOC115960203 | AT4G22330 | AtACER/ATCES1 | ABA response | Unknown/unclassified | T1→T3 | 11 | 0.89 |
| LOC115988690 | AT4G21350 | B80/PUB8 | ABA response | Unknown/unclassified | T2→T3 | 9.1 | 0.8 |
| LOC115956114 | AT3G11130 | AtCHC1/CHC1/HAS1 | water deprivation | Unknown/unclassified | T1→T3 | 9.6 | 0.76 |
| LOC115968434 | AT2G14260 | PAP1/PIP | water deprivation | Unknown/unclassified | T2→T3 | 10.1 | 0.75 |

| | | | | | | | |
|---|---|---|---|---|---|---|---|
| LOC115982354 | AT3G26090 | ATRGS1/RGS1 | ABA response; water deprivation | Unknown/unclassified | T1→T3 | 14.5 | 0.75 |

**Table S5. Drought-associated changes in candidate new TE insertion rates across three levels of read support.** Candidate insertions were classified as single-read calls, all multi-read calls, or multi-read calls retained after cross-leaf re-genotyping. Treatment effects were estimated by comparing plant-level changes between the pre-treatment leaf (T1) and leaves formed during the experiment (mean of T2 and T3), with insertion rates normalised either by sequencing yield (Gb) or by non-TE insertion rates. Drought/control rate ratios and corresponding Welch 95% confidence intervals and *P* values were calculated from plant-level log-transformed changes (n = 4 plants per treatment). A rate ratio of 1 indicates no difference in temporal change between treatments.

| Read-support category | Normalisation | Rate ratio | 95% CI | P |
|---|---|---|---|---|
| Single-read | Per Gb | 0.87 | 0.46-1.62 | 0.569 |
| Single-read | Non-TE calls | 1.01 | 0.88-1.17 | 0.834 |
| Multi-read, all candidates | Per Gb | 1.13 | 0.66-1.92 | 0.592 |
| Multi-read, all candidates | Non-TE calls | 0.85 | 0.56-1.29 | 0.378 |
| Multi-read, re-genotyped | Per Gb | 3.14 | 0.24-41.14 | 0.261 |
| Multi-read, re-genotyped | Non-TE calls | 2.26 | 0.26-19.27 | 0.383 |

**Table S6. Within-leaf allele fractions of multi-read candidate insertions retained after cross-leaf re-genotyping**. For TE and non-TE insertions detected in individual leaves, the total number of candidates, median and 90th-percentile within-leaf allele fractions, and number of candidates with an allele fraction ≥0.25 are reported separately for drought-treated and well-watered plants. Allele fraction was defined as the proportion of reads at the locus supporting the candidate insertion.

| Insertion type | Treatment | Calls | Median allele fraction | 90th percentile | Calls ≥0.25 |
|---|---|---|---|---|---|
| TE | Control | 120 | 0.052 | 0.111 | 0 |
| TE | Drought | 111 | 0.054 | 0.118 | 0 |
| Non-TE | Control | 86 | 0.041 | 0.091 | 0 |
| Non-TE | Drought | 61 | 0.051 | 0.13 | 1 |
